# Informational architecture organizes plasmid genomes

**DOI:** 10.64898/2026.09.01.748738

**Authors:** Yuxuan Zhong, Teng Wang

**Affiliations:** College of Life Sciences and Oceanography, Shenzhen University, Shenzhen 518055, China; Shenzhen University of Advanced Technology, Shenzhen 518107, China; State Key Laboratory of Quantitative Synthetic Biology, Shenzhen Institute of Synthetic Biology, Shenzhen Institutes of Advanced Technology, Chinese Academy of Sciences, Shenzhen, 518055, China

## Abstract

Plasmids are widely viewed as carriers of accessory genes, whereas molecular machineries required to replicate, transcribe and translate genetic information are thought to be encoded primarily by chromosomes. Yet, the recurrent presence of these central dogma-associated genes on plasmids raises a fundamental question of whether these genes merely represent incidental acquisitions or form coordinated informational components that support plasmid function. Here, by analyzing 61,961 plasmids spanning diverse prokaryotic lineages, we show that plasmid-encoded informational genes are not random cargos but form predictable architecture. Using plasmid-borne tRNAs as a model, we demonstrate that their presence, composition and abundance can be accurately inferred from surrounding plasmid gene content and are coupled to the codon demands of plasmid genes. Similar patterns extend to genes associated with replication, transcription, and translation, revealing coordinated informational organization across plasmid genomes. Together, these findings suggest that plasmids do not only encode adaptive functions but also propagate the infrastructure required to execute them, expanding the evolutionary role of horizontal gene transfer.

## Introduction

The flow of genetic information through replication, transcription, and translation defines the molecular logic of life^1^. In prokaryotes, this informational machinery is thought to be encoded primarily by chromosomes, whereas plasmids are typically viewed as dispensable and accessory elements that disseminate adaptive traits such as antibiotic resistance, virulence and metabolic capabilities^2–5^. Yet many plasmids carry genes associated with these central-dogma processes, including transfer RNAs (tRNAs), ribosomal proteins, aminoacyl-tRNA synthetases, and transcription-related factors^6–11^, thereby challenging the well-recognized assumption of microbial genome organization: informational functions belong to chromosomes and adaptive functions to plasmids. More fundamentally, it raises the question of whether these genes are randomly accumulated through gene exchange or are organized into coordinated architectures that support plasmid function^9^.

Distinguishing these possibilities requires moving beyond the mere presence of informational genes to ask whether they are organized within a broader genomic context. Genes acquired opportunistically are expected to occur independently of the context, without functional coupling to neighboring plasmid genes. Conversely, if they contribute to a coordinated plasmid program, they should exhibit predictable relationships with plasmid gene content, expression demands, and other informational components^12,13^. Such predictability would reveal that some plasmids are not simply carriers of individual functions, but structured systems that encode part of the molecular infrastructure required for their own maintenance and expression^4,14^.

Transfer RNAs (tRNAs) provide a powerful lens to examine these two possibilities. As the molecular interface between genetic information and protein synthesis, tRNAs directly connect informational capability with translational demand^15–18^. Despite their essential role in cellular function, tRNA genes are repeatedly found on plasmids across diverse microbial lineages^7,9^. This paradox—an indispensable component of gene expression carried by mobile genetic elements—makes plasmid-borne tRNAs an ideal system for testing whether plasmids encode organized informational architectures. If such architectures exist, plasmid tRNA repertoires should be predictable from the broader context of plasmid genomes.

In this work, we first use plasmid-encoded tRNAs as a model to test whether informational genes are organized within plasmid genomes. Across a comprehensive collection of bacterial and archaeal plasmids, we show that tRNA repertoires are highly structured, predictable from plasmid gene content, and coupled to the translational demands of plasmid genes. This organization further extends to other replication-, transcription-, and translation-associated functions. Together, these results identify predictable informational architecture as an organizing principle of plasmid genomes and indicate that horizontal gene transfer can disseminate not only adaptive functions, but also informational infrastructure coupled with plasmid genetic programs.

## Results

### Distribution pattern of plasmid tRNAs across taxa

To systematically examine plasmid-encoded tRNAs across prokaryotes, we curated 50,720 complete bacterial and archaeal genomes from the NCBI RefSeq database (released before June 2025; Fig. 1A). Each assembly was annotated by the Prokaryotic Genome Annotation Pipeline (PGAP)^19–21^, which provided uniform identification of coding sequences, including tRNAs predicted by tRNAscan-SE^22,23^, and consistent classification of replicons as chromosomes or plasmids, enabling systematic cross-genome comparison of plasmid-borne tRNA genes.

**Figure 1.**
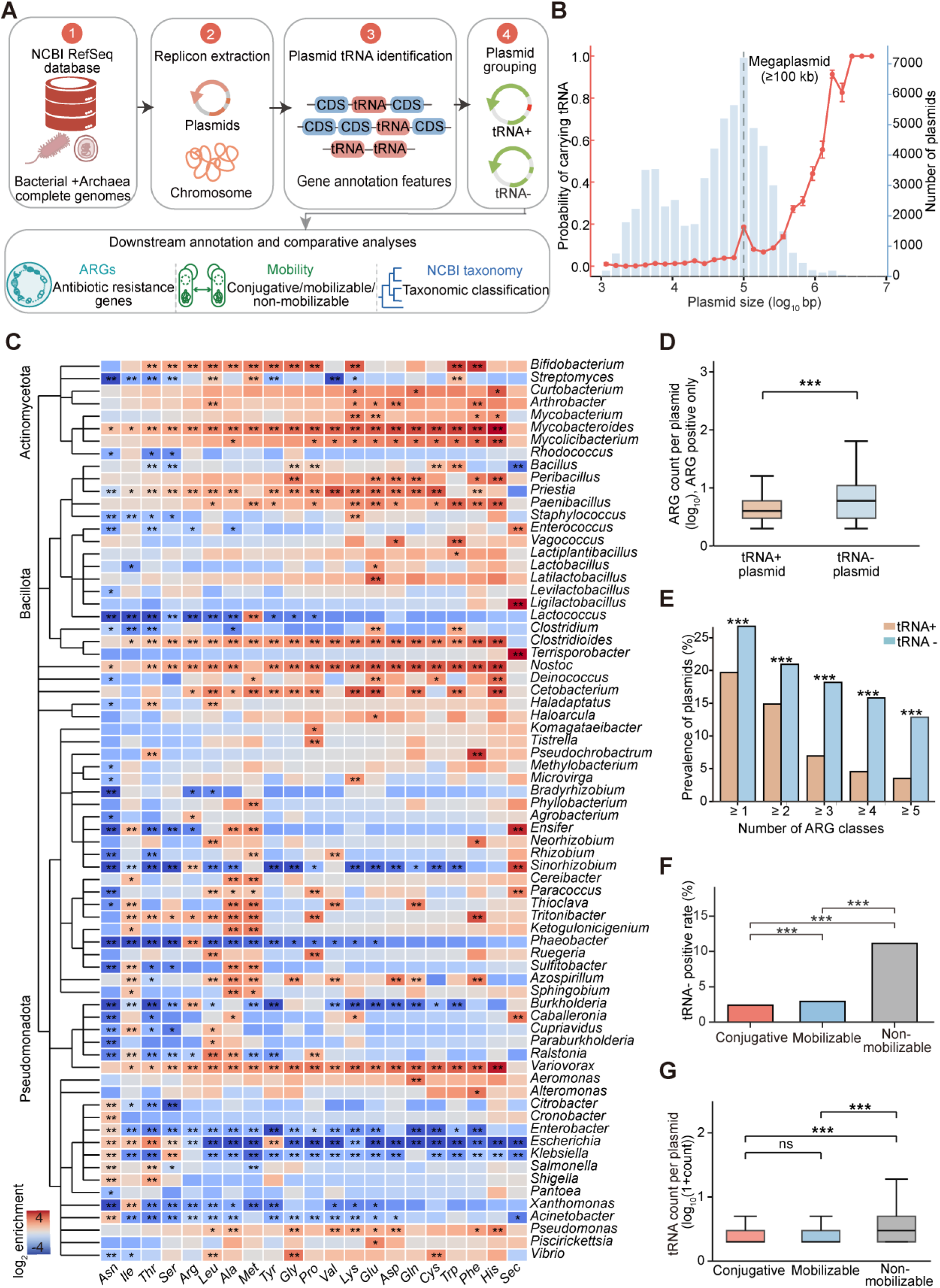
Global landscape of plasmid-encoded tRNAs. (A) Overview of dataset construction and analysis workflow. Complete bacterial and archaeal genomes were retrieved from NCBI RefSeq, plasmid replicons were identified from genome annotations, and plasmid-encoded tRNAs were characterized across genomes. (B) Relationship between plasmid size and tRNA carriage. Blue bars indicate plasmid counts within size bins, and the red curve indicates the fraction of plasmids encoding at least one tRNA. The dashed line marks the 100-kb threshold. (C) Genus-level enrichment landscape of plasmid-encoded tRNA types. Colors represent log2 enrichment relative to the global plasmid background; red and blue indicate enrichment and depletion, respectively. Each genus-tRNA association was tested using a two-sided Fisher’s exact test, with Benjamini-Hochberg adjustment. (D) Distribution of antibiotic-resistance-gene (ARG) counts among ARG-positive plasmids stratified by tRNA carriage. Group comparisons used two-sided Mann–Whitney U test. (E) Prevalence of resistance-class richness among tRNA-positive and tRNA-negative plasmids. At each threshold, the two groups were compared using a two-sided Fisher’s exact test, with Benjamini-Hochberg adjustment across the five tests. (F) Frequency of tRNA carriage across plasmid mobility classes. All three pairwise comparisons were performed using two-sided Fisher’s exact tests with Benjamini-Hochberg adjustment. (G) Number of encoded tRNAs among tRNA-positive plasmids stratified by mobility class. The omnibus comparison used a Kruskal-Wallis test, followed by all three pairwise two-sided Mann-Whitney U tests with Benjamini-Hochberg adjustment. Boxes show the interquartile range (IQR) and median; whiskers extend to the most extreme values within 1.5 × IQR. In D, the asterisk denotes the raw P value; in C and E-G, asterisks denote Benjamini-Hochberg-adjusted q values: ns, P or q >= 0.05; *P or q < 0.05; **P or q < 0.01; ***P or q < 0.001.

From these genomes, we identified 61,961 plasmids, of which 4,125 (6.66%) encoded at least one tRNA gene, suggesting that tRNA carriage is a recurrent rather than exceptional plasmid feature (Fig. 1B). Collectively, plasmid-borne tRNAs cover the full translational coding space, including the whole set of 20 canonical amino acids and selenocysteine^24^. However, this coverage is highly structured, with pronounced preference towards specific tRNA types, particularly tRNA-Asn, tRNA-Ile, and tRNA-Thr. At the phylogenetic level, tRNA-bearing plasmids were widely distributed but exhibited marked lineage-specific enrichment patterns (Fig. 1C). For example, *Mycobacteroides* showed enrichment for nearly all plasmid-encoded tRNA types, whereas *Escherichia* displayed depletion for most (see Methods for details). Notably, 91.93% of tRNA-positive plasmids encoded tRNA types that were also present in the host chromosome. Among the remaining 8.07% carrying at least one host-absent tRNA type, 87.39% were megaplasmids (≥100 kb)^25,26^.

Having established the widespread yet non-random presence of plasmid-encoded tRNAs, we next asked which genomic features shape their distributions. tRNA-positive plasmids were substantially larger than tRNA-negative plasmids (Fig. S1A) and showed modest but consistent increases in GC content (Fig. S1B) and coding density (Fig. S1C). Notably, despite their expanded coding capacity, tRNA-positive plasmids were depleted in antibiotic resistance genes (ARGs)^27^ (Fig. 1D; Fig. S1D) and associated with lower multidrug resistance burden (Fig. 1E; Fig. S1E). This trend extends to plasmid mobility^28^: tRNA genes were rare in conjugative and mobilizable plasmids, yet markedly enriched in non-mobilizable plasmids (Fig. 1F; Fig. S1F), which also encoded larger tRNA repertoires (Fig. 1G; Fig. S1G). Taken together, these results revealed the tight association between tRNA-carriage and diverse plasmid traits.

### Plasmid gene content predicts tRNA carriage and composition

The structured distribution of plasmid-encoded tRNAs pointed to their integration into the broader functional organization of plasmid genomes. Thus, tRNA carriage might be predictable from the non-tRNA genes residing on the same plasmid. To test this hypothesis, we represented each plasmid by the abundances of 29,267 non-tRNA gene product descriptors extracted from PGAP annotations, and trained separate models to predict either tRNA presence by classification or tRNA abundance by regression (Fig. 2A). All seven classifiers underwent hyperparameter optimization using 50 candidate configurations per classifier and randomization, evaluated exclusively on an internal validation subset of each training partition using AUPRC as the selection criterion (Fig. S2A). Using only non-tRNA gene content, LightGBM^29^ was selected as the downstream modelling framework, achieving a mean test AUPRC of 0.9042 ± 0.0118 with consistently high AUROC, precision, recall and F1 scores across three independent 80:20 train–test splits (Fig. 2B; Fig. S2B; see Methods for more details), and was retained as the general framework owing to its unified support for classification, regression and gain-based feature interpretation, together with lower training time and memory requirements in this high-dimensional sparse-data setting.

**Figure 2.**
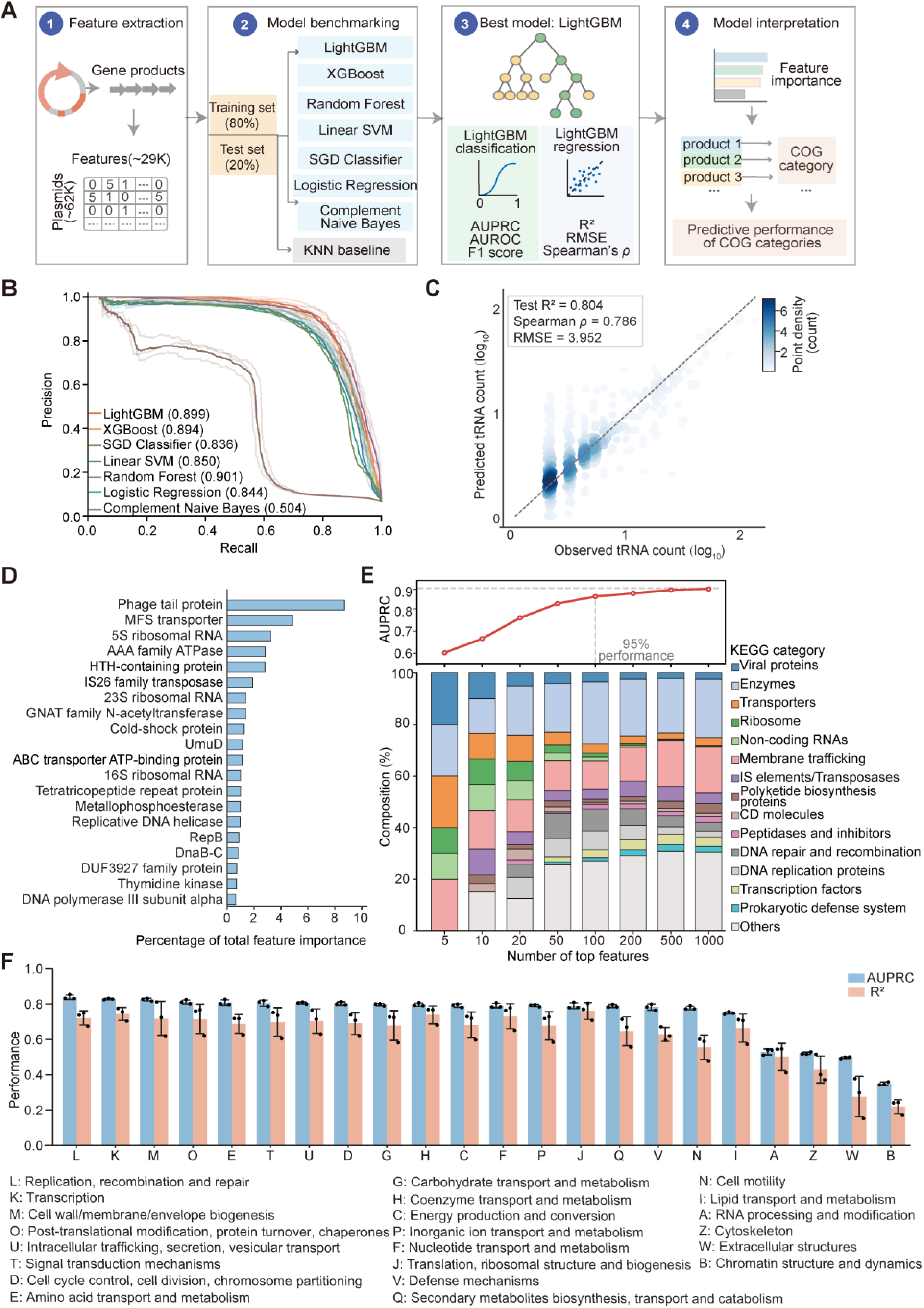
Plasmid tRNA carriage and abundance is predictable from surrounding plasmid gene content. (A) Machine-learning workflow. Plasmid gene-product annotations were converted into a high-dimensional feature matrix after removal of tRNA genes. The resulting non-tRNA features were used to predict binary tRNA carriage, total tRNA abundance, and tRNA type-specific abundances. (B) Precision-recall curves for seven classifiers evaluated on held-out test partitions after validation-only hyperparameter selection. (C) LightGBM regression of total plasmid tRNA count. Observed and predicted values are shown on a log10 scale. (D) Feature importance for classification. Top LightGBM features were ranked by gain-based importance. (E) Top-K feature sufficiency and functional composition analysis. The upper curve shows the AUPRC obtained using the top-K ranked features. The lower stacked bars show the functional composition of the top-ranked features. (F) Predictive performance of individual COG/NOG categories for binary classification and regression. Machine-learning summaries show the mean across three independent random 80:20 train-test splits, error bars show standard deviation, and black points show the replicate values. Classification was evaluated by AUPRC and regression by R² on held-out test partitions.

This predictability extended beyond tRNA presence to tRNA repertoire composition. Plasmid gene content accurately predicted total tRNA abundance (R² = 0.8040 ± 0.0605; RMSE = 3.9517 ± 0.5337; Spearman’s *ρ* = 0.7864 ± 0.0202; MAE = 1.2599 ± 0.1516; Fig. 2C) and individual tRNA species, with variable performance across types (Fig. S3A and B). For tRNA-Sec, all positive plasmids carried a single copy, precluding variance-based regression metrics. In comparison, similarity-based nearest-neighbor models performed substantially worse for both classification and regression (Fig. S4A and B), indicating that predictive power stems from coordinated functional organization rather than gene-content similarity alone. Models using only host-chromosomal gene content performed poorly (mean AUPRC = 0.3214; mean R² = 0.1069; Fig. S4C), indicating that the predictive signal resides primarily in the plasmid’s own gene repertoire rather than in its host background.

To assess whether prediction reflected genuine plasmid gene–tRNA coupling rather than shared evolutionary history, we evaluated model performance under increasingly stringent partitioning strategies (Fig. S5A and B). Performance was preserved in genome-blocked evaluations, where all plasmids from the same host genome were restricted to a single partition, demonstrating generalization beyond individual genome backgrounds. As expected, performance decreased under species- and genus-level blocking, consistent with lineage-specific organization of plasmid gene repertoires. Nevertheless, predictive performance remained substantially above baseline, indicating that the observed signal was not solely attributable to phylogenetic similarity.

We next examined whether prediction could be explained by plasmid size or annotation characteristics. Although larger plasmids showed stronger predictability, models retained robust performance within individual plasmid size classes (Fig. S5C). Cross-size transfer experiments further demonstrated that predictive signals were not solely driven by plasmid size, although generalization was asymmetric: models trained on large plasmids retained substantial predictive power when applied to small plasmids, whereas models trained on small plasmids showed reduced performance on large plasmids (Fig. S5D). To further test whether prediction was driven by plasmid size, we repeated the analysis using length-normalized product densities (per 100 kb). Classification performance remained essentially unchanged, whereas abundance prediction showed only a modest decline (Fig. S5E and F). Performance also remained robust across major host phyla (Fig. S6A and B), and removal of the standardized hypothetical-protein feature had minimal impact on either classification or abundance prediction (Fig. S7), excluding annotation-dependent artifacts as a major source of prediction.

To evaluate temporal generalization, we applied models trained exclusively on the original RefSeq cohort to an independent set of plasmid genomes released after model development. Without retraining, feature modification, or threshold recalibration, the models retained high performance (mean AUPRC = 0.9108; mean R² = 0.8441; Fig. S8). These results demonstrate that the association between plasmid gene repertoires and tRNA organization is not restricted to the original training cohort but represents a transferable genomic signature.

To assess whether the observed predictive architecture was sensitive to potential replicon misclassification, we independently reclassified tRNA-bearing replicons using PlasFlow^30^. According to PlasFlow classification, 61.2% of RefSeq-annotated plasmids were assigned plasmid-like, whereas 14.9% were classified as chromosome-like and 23.9% remained unresolved (Fig. S9A). We then restricted the existing held-out test-set predictions to PlasFlow-supported plasmid-like replicons and recalculated abundance-prediction metrics without retraining the models or modifying the feature space. tRNA abundance prediction remained substantial within this subset (Fig. S9B), indicating that the predictive signal was not driven solely by potentially misclassified replicons.

### Functional architecture underlying plasmid tRNA predictability

Which plasmid functions encode this predictive information? Feature-importance analysis of the LightGBM classifier revealed that prediction was concentrated in a small subset of product descriptors, with the top 100 features retaining 95.3% of full-model performance (Fig. 2D, E; see Methods). These features were enriched for replication and genome maintenance, phage-associated functions, transposition, transport, stress responses, and translation-related processes^31^. The regression model revealed a more pronounced contribution from translation-associated functions, with the strongest predictors including SsrA-binding protein SmpB, ribosomal RNAs, elongation factor Tu, ribosomal proteins, and peptidyl-tRNA hydrolase (Fig. S10A). Similar to classification, regression performance was largely captured by a limited feature set, with the top 100 features retaining >95% of the full-model R² (Fig. S10B). Early-ranked regression features were dominated by ribosomal and translation-related functions, whereas additional features progressively incorporated broader processes, including ribosome biogenesis, membrane transport, enzymatic functions, and DNA repair and recombination.

To determine whether these signals reflected broader functional organization rather than individual predictive features, we trained independent models using genes assigned to individual COG categories (see Methods for details). Consistent with feature-importance analysis, COG-based models identified replication, recombination and repair (L) as the strongest predictive category, followed by transcription (K), cell envelope biogenesis (M), protein turnover (O), amino acid transport and metabolism (E), and signal transduction (T) (Fig. 2F)^32,33^. These results indicate that plasmid tRNA repertoires are encoded within a distributed functional architecture that links translational capacity with plasmid maintenance, regulation, and adaptation.

Finally, we examined whether this architecture was shared across individual tRNA types. Feature-importance profiles from classification and regression models for each of the 21 tRNA types revealed both conserved and specialized determinants (Fig. S11; S12). Several highly ranked product descriptors recurred across multiple tRNA models, whereas others showed selective associations with specific tRNA types. Moreover, classification and regression models captured overlapping but distinct feature landscapes, indicating that the genomic determinants of tRNA presence differ from those governing copy-number variation.

### Plasmid-encoded tRNAs are aligned with codon usage of plasmid genes

A defining feature of an informational architecture is that its components should be integrated with the genetic programs they support. We therefore asked whether plasmid-encoded tRNAs are embedded within plasmid functional organization by testing whether plasmid genes preferentially use codons decoded by their associated tRNA repertoires. To quantify this relationship, we developed a codon compensation index (CCI), which measures enrichment of codons decoded by plasmid-borne tRNAs relative to the expectation under unbiased synonymous codon usage (see Methods)^16–18^. A CCI greater than 1 indicates preferential use of codons supported by plasmid-encoded tRNAs (Fig. 3A).

**Figure 3.**
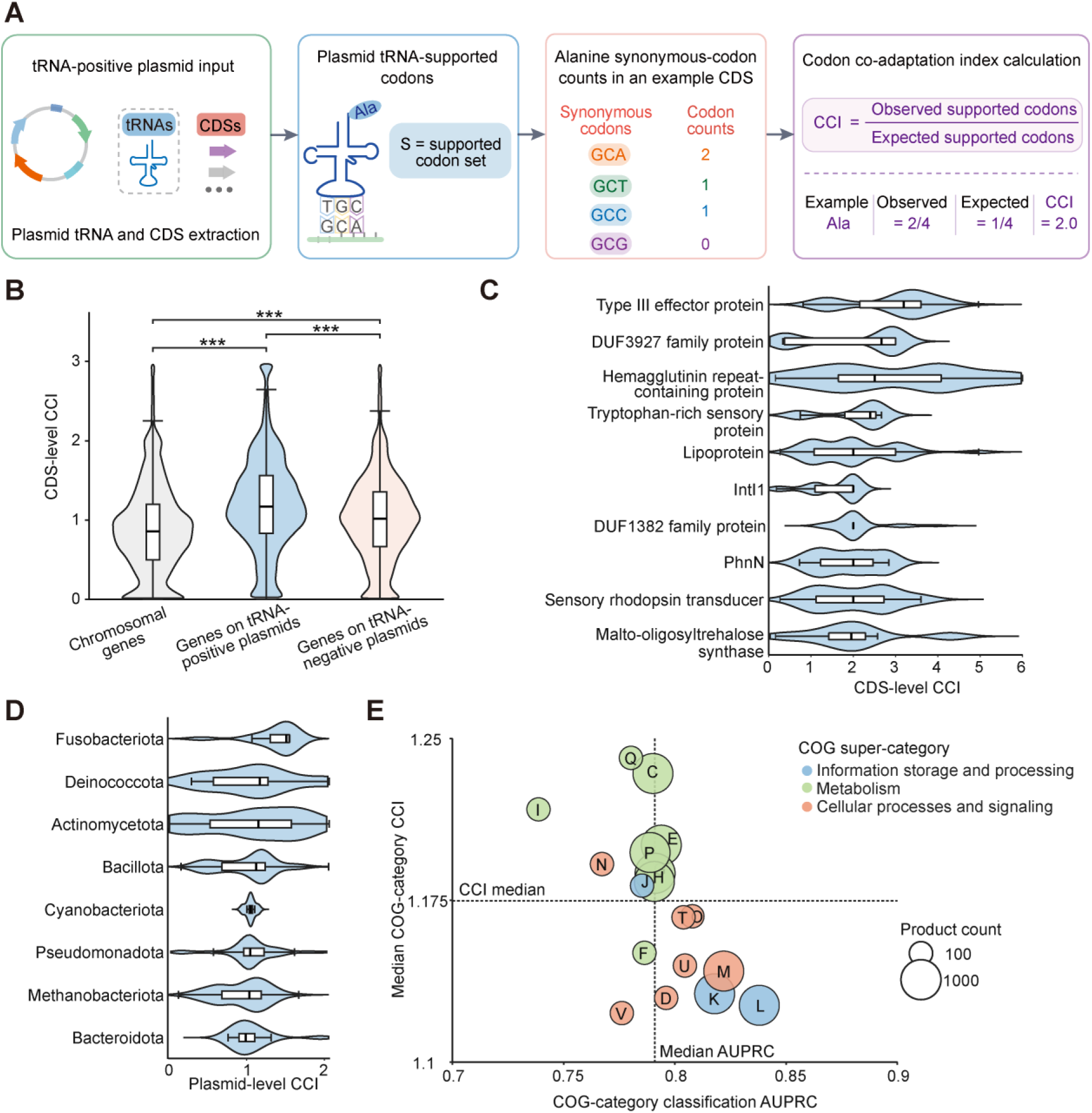
Codon compensation index (CCI) links plasmid-encoded tRNAs to codon usage of plasmid genes. (A) Schematic definition of plasmid-encoded tRNA supply, codon demand, and codon compensation index (CCI). CCI quantifies enrichment of codons supported by plasmid-encoded tRNAs relative to synonymous expectation. (B) CDS-level CCI distributions for chromosomal, tRNA+ plasmid, and tRNA− plasmid genes. CDS-level CCI is the CCI of each individual valid CDS, calculated using the genome-pooled plasmid-tRNA repertoire. Global differences were assessed by Kruskal–Wallis test followed by pairwise two-sided Mann–Whitney U tests with Benjamini–Hochberg correction. *q < 0.05; **q < 0.01; ***q < 0.001. (C) Product-level aggregation of CCI values. For each normalized product descriptor, CCI values were summarized by the median across CDSs; only products represented by more than 50 genes were retained. (D) Plasmid-level CCI distributions across host phyla. Plasmid-level CCI was calculated directly from pooled codon counts across all valid CDSs on each tRNA-positive plasmid using that plasmid’s own tRNA repertoire. Only phyla with at least 10 plasmids are shown. (E) Relationship between COG-category median product-level CCI and classification performance of COG-restricted models. Each point represents one COG category. The x-axis shows the mean classification AUPRC across three independent 80:20 splits for LightGBM models trained using only features assigned to that COG category, and the y-axis shows the median product-level CCI of products assigned to the same category. Point size indicates the number of products assigned to each COG category, and colors indicate COG functional superclasses. Only categories represented by more than 50 products were retained. Dashed lines indicate the median AUPRC and median CCI across the retained COG categories.

Using the pooled plasmid-encoded tRNA repertoire within each genome, we calculated CCI values for all coding sequences in genomes carrying both tRNA-positive and tRNA-negative plasmids (Fig. 3B; Fig. S13A and B). At the individual CDS level, median CCI values were 1.143 for genes on tRNA- positive plasmids, 1.012 for genes on tRNA-negative plasmids, and 0.867 for chromosomal genes, with 60.7%, 50.5%, and 37.4% of CDSs above 1, respectively. All these differences were statistically significant (Kruskal-Wallis P < 2.2 × 10^-308^; all pairwise Mann-Whitney U tests with BH-adjusted q < 2.2 × 10^-308^). Together, these results show that CDSs carried by tRNA-bearing plasmids preferentially use codons supported by their plasmid-derived tRNAs, suggesting that plasmid-encoded tRNAs are integrated with plasmid gene-expression demands rather than representing neutral genetic cargo.

To identify plasmid functions most closely associated with plasmid-derived translational resources, we aggregated CCI values by annotated gene products. Several recurrent plasmid-encoded proteins, including type III effector protein, hemagglutinin repeat-containing protein, and tryptophan-rich sensory protein, showed exceptionally high codon compensation, all with median CCI values exceeding 2.0 (Fig. 3C). These genes therefore represent candidate translational dependents whose expression is particularly aligned with plasmid-borne tRNA repertoires.

Codon compensation also exhibited phylogenetic structure. Treating each plasmid as an integrated coding unit, we calculated plasmid-level CCI by aggregating all encoded CDSs within individual plasmids. Plasmids from *Fusobacteriota*, *Deinococcota*, *Actinomycetota*, and *Bacillota* displayed the highest compensation scores (Fig. 3D), indicating lineage-specific differences in the extent to which plasmid gene repertoires are aligned with their encoded translational resources.

Integrating CCI with LightGBM performance revealed a two-layer organization of plasmid informational architecture (Fig. 3E; Fig. S14). Replication (L), transcription (K), and envelope biogenesis (M) were strong predictors of tRNA carriage but showed weaker codon-level coupling, whereas lipid metabolism (I), secondary metabolism (Q), and related functions showed the opposite pattern. Regression analyses revealed a similar but non-identical organization (Fig. S14). Thus, the genomic context that predicts plasmid-borne tRNAs is partly distinct from the functions that directly exploit them, supporting a distributed architecture in which plasmid tRNAs serve as shared translational resources rather than gene-specific adaptations.

### Extension to broader central-dogma modules

The structured organization observed for plasmid-encoded tRNAs raised the question of whether predictability extends to broader informational functions. We therefore examined plasmid genes involved in the three core processes of the central dogma: replication, transcription, and translation (Fig. 4A; see Methods for details). These modules showed distinct prevalence patterns across plasmids: replication-associated genes were widespread (83.5%), transcription-associated genes were common (58.4%), whereas translation-associated genes are comparatively rare (13.6%). Thus, plasmids frequently encode partial informational modules, with different levels of investment across different central dogma processes. Moreover, replication, transcription, and translation submodules exhibited strong lineage-specific distributions across host taxa (Fig. 4B; Fig. S15), indicating that plasmid informational functions are non-randomly organized across microbial diversity.

**Figure 4.**
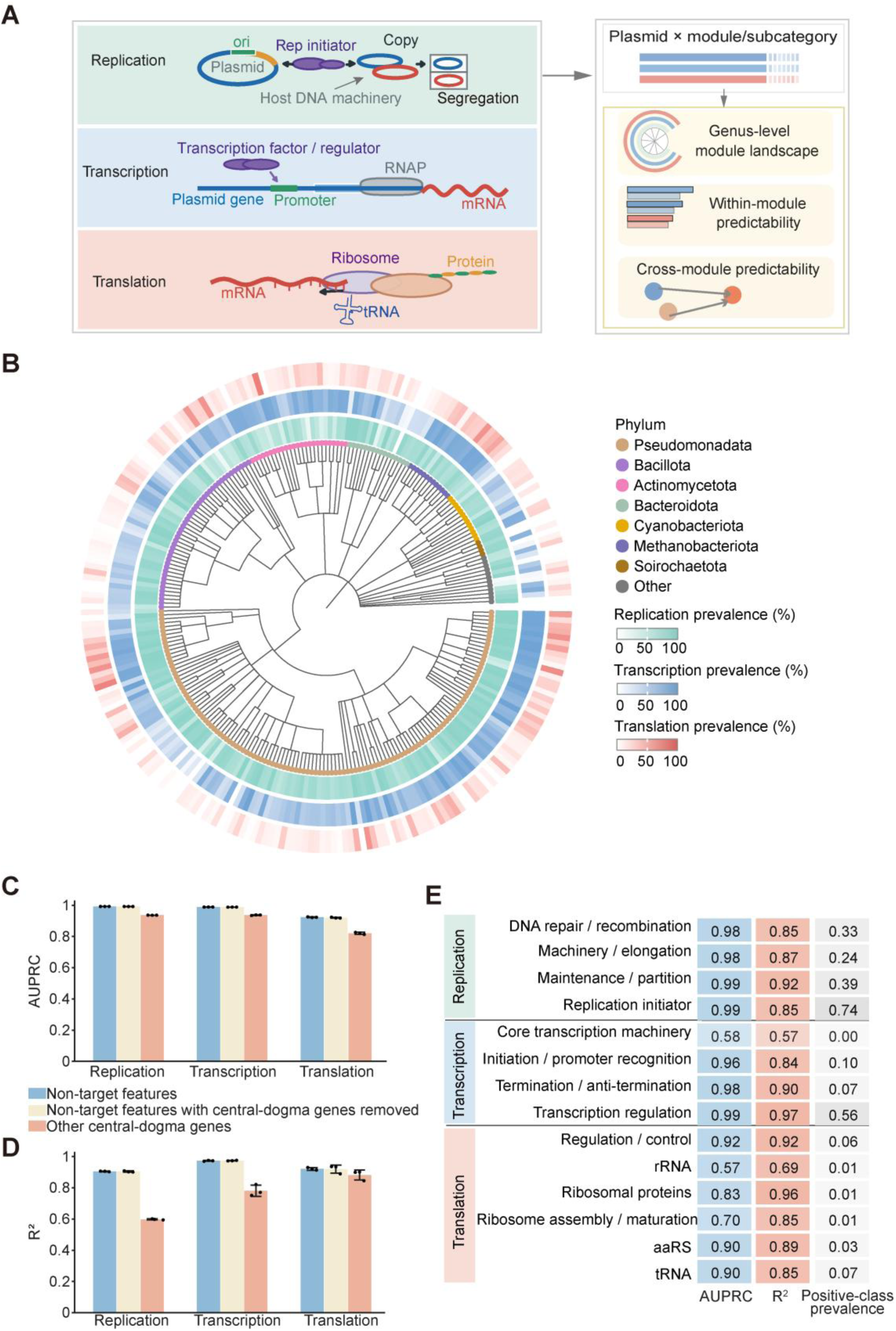
Central-dogma modules exhibit coordinated plasmid architecture. (A) Workflow for manual curation of replication, transcription, and translation modules and subcategories. Target-module features were removed prior to prediction. (B) Taxonomic distribution of central-dogma modules across plasmids. For each host phylum, prevalence was calculated as the fraction of plasmids carrying at least one gene assigned to the indicated module or subcategory among all plasmids from that phylum. (C) Classification performance (AUPRC) for predicting the presence or absence of replication, transcription, and translation modules under three feature settings: all non-target features, non-central-dogma background features, and other central-dogma modules only. Models were evaluated across three target-stratified random 80:20 train-test replicates; each replicate represents an independent split of plasmids into training and held-out test sets. Bars show mean held-out AUPRC, error bars show standard deviations, and black points show raw replicate values. (D) Regression performance (R²) for predicting module copy numbers under the same feature settings as in (C), summarized across the same three train-test replicates. (E) Subcategory-level predictive performance using non-target features, with classification evaluated by AUPRC and regression by R². Positive-class prevalence is shown as a reference baseline and was calculated as the fraction of held-out test plasmids positive for each target subcategory in each replicate.

To determine whether these functions are embedded within broader plasmid architectures, we trained predictive models after removing features belonging to the target module. All three central-dogma modules were highly predictable from the remaining plasmid gene repertoire (Fig. 4C). Replication and transcription showed near-perfect predictability (AUPRC = 0.992 and 0.988, respectively), whereas translation remained highly predictable despite its lower prevalence (AUPRC = 0.921). Regression models similarly recovered module abundance with high accuracy (R² = 0.900– 0.976) (Fig. 4D). Replication, transcription, and translation modules also retained predictive power for one another, although with reduced accuracy (Fig. 4C and D). Importantly, removing all central-dogma-associated features had little effect on performance, demonstrating that these modules are encoded within the broader functional organization of plasmids rather than inferred simply from the co-occurrence of other informational genes (Fig. 4C and D).

This organization extended to finer functional units: replication initiators, DNA repair systems, transcription regulators, aminoacyl-tRNA synthetases, and tRNAs remained highly predictable from the remaining plasmid gene repertoire (Fig. 4E). Independent COG-based analyses yielded similar results: after exclusion of the corresponding target categories, replication (L), transcription (K), and translation (J) functions remained highly predictable from non-target plasmid features (Fig. S16). Together, these findings generalize the concept of informational architecture beyond plasmid-encoded tRNAs, revealing replication, transcription, and translation functions as coordinated and predictable components of mobile genomes.

## Discussion

A prevailing view in microbial evolution is that chromosomes encode the informational machinery required for gene expression, whereas plasmids primarily serve as vehicles for adaptive traits^13,34^ . Our findings challenge this dichotomy. Across more than 60,000 plasmids, tRNAs and other central-dogma-associated functions are not isolated genetic additions, but components of predictable genomic architectures. Their presence, abundance, and functional composition can be inferred from surrounding plasmid gene content, revealing that many plasmids encode not only biological functions, but also elements of the informational capacity required to express them.

The organization of plasmid-encoded tRNAs provides insight into how such architectures are assembled. Genes with greatest predictive power for tRNA carriage are involved in functions of replication, regulation, and genome maintenance, whereas genes most dependent on plasmid-derived translational resources occupy a distinct functional space (Fig. 3E). This separation suggests that plasmid tRNAs are not dedicated to individual targets, but act as shared resources integrated into broader plasmid programs. Translational provisioning therefore emerges as a property of plasmid systems rather than individual genes.

This principle extends beyond tRNAs. Replication, transcription, and translation modules remain predictable even after removing the corresponding informational genes from model inputs, demonstrating that central-dogma functions are embedded within the broader organization of plasmid genomes (Fig. 4C and D). Rather than accumulating informational genes independently, plasmids appear to acquire coordinated modules that couple adaptive capacity with the molecular infrastructure required for its expression. Such coupling may become increasingly important as plasmids expand in size and functional complexity.

These findings suggest a broader revision of how horizontal gene transfer is conceptualized. Mobile genetic elements are typically viewed as vehicles that disseminate individual genes^35,36^; however, our results indicate that they can also propagate higher-order informational architectures that link genetic content with expression capacity. In this view, horizontal transfer does not simply redistribute biological functions—it can reshape the molecular systems through which those functions are realized.

Informational architecture therefore represents a previously underappreciated level of genome organization, positioned between individual genes and complete genomes. Similar principles may extend beyond plasmids to other mobile genetic elements and may provide a framework for understanding how complex genetic systems evolve, spread, and become functionally integrated^4,36–38^. Beyond evolutionary biology, identifying the rules governing such architectures may also inform the design of synthetic vectors^39,40^, where predictable gene expression across diverse hosts remains a central challenge.

## Methods

### Genome dataset and plasmid annotation

Complete prokaryotic genomes were retrieved from NCBI RefSeq as of June 2025. Only assemblies annotated by the NCBI Prokaryotic Genome Annotation Pipeline (PGAP)^21^, designated as “Complete Genome”, and not flagged as atypical were retained. The final dataset contained 50,720 genomes, including 50,092 bacterial and 628 archaeal assemblies.

GenBank flat files (GBFF) were parsed to extract replicon-level and CDS-level annotations. Replicons were classified as chromosomes or plasmids based on source-feature qualifiers, with record metadata used as secondary evidence when necessary. For each replicon, we extracted replicon size, nucleotide sequence, CDS product annotations, amino acid sequences, and annotated tRNA features. Plasmid-related data were aggregated into a dataset containing 61,961 plasmids.

Plasmid-encoded tRNA genes were directly identified from annotated tRNA features in GBFF files. For each plasmid, we recorded the tRNA amino acid specificity and copy number. Plasmids were classified as tRNA-positive if at least one tRNA gene was present, and tRNA-negative otherwise, yielding 4,125 tRNA-positive and 57,836 tRNA-negative plasmids, respectively. Comparisons were performed between tRNA-positive and tRNA-negative plasmids, with differences in plasmid size, GC content, and coding density evaluated using two-sided Mann–Whitney U tests^41^. Coding density was calculated from the original GBFF annotations as the number of plasmid nucleotides covered by at least one annotated CDS divided by the plasmid sequence length. To examine size-dependent effects, plasmids were additionally binned on log scale, and tRNA carriage probabilities were computed across size intervals, with ≥100 kb defined as the threshold for megaplasmids^25^.

To independently evaluate the sequence identity of tRNA-bearing plasmids, plasmid nucleotide sequences were classified using PlasFlow, a neural network-based plasmid sequence classifier^30^. PlasFlow predictions were performed on all RefSeq-annotated tRNA-positive plasmids using default parameters. Each sequence was assigned to the predicted category with the highest confidence, including plasmid-like, chromosome-like, or unclassified. PlasFlow classifications were used to assess whether model performance and major biological patterns were preserved among PlasFlow-supported plasmid sequences.

### Taxonomic assignment and genus-level enrichment analysis

Assembly accessions were joined to the NCBI RefSeq Assembly Summary file to obtain NCBI Taxonomy IDs and organism names. For records with valid Taxonomy IDs, taxonomic lineages were retrieved using the NCBITaxa module^42^ and standardized at the superkingdom, phylum, class, order, family, genus, and species levels. Of the 23,248 target assemblies that contained annotated plasmid replicons, 23,090 were matched to the current taxonomy table and included in taxonomy-dependent analyses, whereas the remaining 158 lacked a matching taxonomy record and were excluded from those analyses. Records with missing genus assignments or genus labels annotated as unknown or uncultured were further excluded from genus-level analyses.

For genus-level enrichment analysis, genera represented by fewer than five tRNA-positive plasmids were excluded to minimize sampling bias from sparsely represented taxa, resulting in 83 retained genera and 3,670 plasmids. For each plasmid, plasmid-encoded tRNA features were converted to binary variables based on the presence or absence of each tRNA type. For each genus–tRNA pair, enrichment or depletion was assessed using Fisher’s exact test on 2×2 contingency tables comparing plasmids from the focal genus with plasmids from all other genera, and plasmid-encoded tRNA carriers with non-carriers. For each focal genus and tRNA type, a denotes the number of plasmids carrying the tRNA in the focal genus, b the number of plasmids not carrying it in the focal genus, c the number of plasmids carrying it in all other retained genera, and d the number of plasmids not carrying it in all other retained genera, with n_in = a + b and n_out = c + d. P values were adjusted using the Benjamini– Hochberg procedure. Enrichment was quantified as the log₂ ratio of tRNA-type prevalence in the focal genus to that in the remaining genera, using 0.5 pseudocounts: (*a* + 0.5)/(*n*_in_ + 1) and (*c* + 0.5)/ (*n*_out_ + 1) . Associations with an adjusted P < 0.05 and llog₂ enrichmentl > 1 were considered statistically significant and retained for further interpretation.

### Antibiotic resistance gene analysis

Antibiotic resistance genes (ARGs) were annotated using the Resistance Gene Identifier against the Comprehensive Antibiotic Resistance Database (CARD)^27^. ARG annotations were aggregated by plasmid to obtain the total number of ARG hits per plasmid. ARG analyses included 61,958 plasmids with available ARG annotation data; three plasmids without ARG results were excluded. Plasmids containing at least one ARG hit were classified as ARG-positive. The proportion of ARG-positive plasmids was compared between tRNA-positive and tRNA-negative plasmids using a two-sided Fisher’s exact test^43^. Among ARG-positive plasmids, the distributions of total ARG counts per plasmid were compared between the two tRNA groups using a two-sided Mann–Whitney U test on untransformed counts. For visualization only, counts were transformed as log10(1 + count). The same comparison was repeated separately for plasmids <100 kb and ≥100 kb, and P values were adjusted using the Benjamini-Hochberg procedure.

Resistance-class richness was defined as the number of distinct antibiotic resistance classes on each plasmid. Plasmids were further categorized according to whether they carried resistance determinants from at least one, two, three, four, or five resistance classes. Multidrug resistance (MDR) was defined as the presence of resistance determinants from three or more antibiotic classes. For each threshold, the prevalence of resistance-class carriage was compared between tRNA-positive and tRNA-negative plasmids using separate two-sided Fisher’s exact tests; the five P values were adjusted jointly using the Benjamini-Hochberg procedure^44^. MDR prevalence was compared between the two tRNA groups separately within the <100 kb and ≥100 kb strata using two-sided Fisher’s exact tests, followed by Benjamini-Hochberg adjustment.

### Plasmid mobility classification

Plasmid mobility classes were assigned using MOB-suite, which classified plasmids as conjugative, mobilizable, or non-mobilizable based on mobility-associated features^28^. tRNA prevalence was calculated within each mobility class as the proportion of plasmids carrying at least one tRNA gene. All three pairwise differences in prevalence were tested using two-sided Fisher’s exact tests, and the three P values were adjusted using the Benjamini-Hochberg procedure. For tRNA abundance analysis, only tRNA-positive plasmids were included. Raw tRNA counts per plasmid were first compared among the three mobility classes using a Kruskal-Wallis test^45^, followed by all three pairwise two-sided Mann-Whitney U tests with Benjamini-Hochberg adjustment. Counts were transformed as log10(1 + count) for visualization only. The prevalence and abundance analyses were also repeated separately for plasmids <100 kb and ≥100 kb; within each size stratum, the three pairwise P values were adjusted as one Benjamini-Hochberg family. These analyses evaluated whether plasmid mobility was associated with the prevalence and abundance of plasmid-encoded tRNAs.

### Feature engineering and machine-learning-based classification

Gene product descriptors were extracted from all plasmids and converted into a plasmid-by-gene feature matrix. Unique product names were standardized and encoded as distinct features, with feature values corresponding to gene counts per plasmid. To prevent data leakage, all tRNA genes were removed from the feature set prior to model construction. The final dataset comprised 61,961 plasmids represented by 29,267 non-tRNA gene features.

To predict plasmid tRNA carriage, we evaluated seven machine-learning classifiers: LightGBM, XGBoost^46^, logistic regression^47^, linear support vector machine^48^, random forest^49^, stochastic gradient descent (SGD)^50^, and Complement Naive Bayes^51^. Models were trained using 80:20 train–test splits and assessed across repeated randomizations. Performance was evaluated using area under the precision– recall curve (AUPRC), area under the receiver operating characteristic curve (AUROC), precision, recall, and F1 score, with AUPRC used as the primary metric because of class imbalance. For each classifier and random seed (42, 43, 44), 50 hyperparameter^52^ configurations were sampled from predefined search spaces and evaluated by AUPRC on an internal validation subset (derived from an additional 80:20 split of the training data). The selected model was then refitted on the complete development set and evaluated on the held-out test set, which was not used for selection. The LightGBM model was optimized using a hyperparameter search. The number of trees was evaluated at {300, 600, 900, 1,200}. The learning rate was sampled from a log-uniform distribution over [0.01, 0.10]. The number of leaves was chosen from {31, 63, 95, 127}, and maximum tree depth was set to either unlimited, 8, 12, or 16. The minimum number of samples per child node was tested at {5, 10, 20, 40, 80}. Row and column subsampling ratios were searched uniformly over [0.7, 1.0] and [0.6, 1.0], respectively. Finally, L1 and L2 regularization parameters were sampled from log-uniform distributions over [10^-4^, 1] and [10⁻⁴, 10], respectively. XGBoost candidates varied the same tree number and learning-rate ranges together with maximum depth, minimum child weight, row and column subsampling, gamma and L1/L2 regularization. Logistic regression and linear SVM candidates varied C, penalty or loss, and class weighting; Random Forest candidates varied tree number, depth, feature subsampling, minimum split size, minimum leaf size and class weighting; SGD candidates varied loss, penalty, alpha, elastic-net mixing and class weighting; and Complement Naive Bayes candidates varied alpha and normalization.

Random forest achieved the highest mean validation AUPRC (0.8992 ± 0.0225), closely followed by LightGBM (0.8982 ± 0.0180), with mean test AUPRCs of 0.9064 ± 0.0211 and 0.9004 ± 0.0151, respectively. Given the small difference and LightGBM’s unified framework for classification, regression, and feature importance, LightGBM was selected for downstream analyses. To obtain a single configuration across seeds, we evaluated the three run-specific winners on all validation splits and selected the one with the highest mean AUPRC. Seeds thus affected only data partitions and random state. Final performance (mean ± SD of AUPRC, AUROC, precision, recall, and F1) was assessed on the three held-out test partitions.

To assess whether model performance could be attributed to simple similarity-based retrieval, we evaluated 1-nearest-neighbor (1-NN) and 5-nearest-neighbor (5-NN) classifiers as baseline approaches^53^. Nearest neighbors were identified using cosine similarity in the non-tRNA gene-feature space. Stratified 80:20 train–test splits were generated using three independent random seeds. For each test plasmid, the 1-NN classifier assigned the tRNA-carriage label of the most similar training plasmid, whereas the 5-NN classifier predicted the proportion of tRNA-positive plasmids among the five nearest training neighbors using uniform weighting.

### Prediction of plasmid-borne tRNA abundance by machine-learning-based regression

Quantitative prediction of plasmid tRNA repertoires was performed using LightGBM regression. Models were trained using 80:20 train–test splits and assessed across repeated randomizations. Performance was evaluated using the coefficient of determination (R²), root mean squared error (RMSE), mean absolute error (MAE), and Spearman correlation between observed and predicted values, with RMSE used as the primary metric for hyperparameter selection. For each random seed (42, 43, 44), 50 hyperparameter configurations were sampled from predefined search spaces and evaluated by RMSE on an internal validation subset (derived from an additional 80:20 split of the training data). The selected model was then refitted on the complete development set and evaluated on the held-out test set, which was not used for selection. To obtain a single configuration across seeds, the three run-specific winners were evaluated across all three validation splits and the configuration with the lowest mean RMSE was fixed. Seeds thus affected only data partitions and random state. Final performance (mean ± SD of R², RMSE, MAE, and Spearman correlation) was assessed on the three held-out test partitions.

### Model robustness and regression analyses

Model robustness was evaluated using progressively more stringent data-partitioning strategies, including random, genome-blocked, species-blocked and genus-blocked splits. In blocked evaluations, plasmids originating from the same genome or host taxonomic group were assigned exclusively to either the training or test set, thereby testing generalization beyond closely related plasmids and host lineages. To assess the contribution of plasmid size to prediction, performance was evaluated across size-stratified subsets and cross-size transfer experiments, in which models trained on small plasmids (<100 kb) were tested on large plasmids (≥100 kb), and vice versa.

To further evaluate whether prediction could be explained by the trivial increase in absolute gene counts with plasmid size, we normalized each product count by plasmid length and repeated the analyses using product densities per 100 kb: 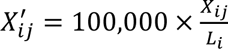, where *X_ij_* is the original product count and *L_i_* is the plasmid length in base pairs. The normalized matrix was evaluated with the same locked LightGBM configuration and identical train–test partitions as the original analysis.

### Host-chromosome baseline

To determine whether plasmid tRNA content could be predicted from host chromosomal genes, non-tRNA product descriptors were extracted from non-plasmid replicons within the corresponding RefSeq assemblies and converted into product-count feature vectors. Standardized non-tRNA product descriptors were collected from chromosome replicons of 23,248 eligible Assembly_IDs, yielding 51,615 distinct chromosome-derived product features. For each Assembly_ID, copy numbers of each product were summed across all chromosome replicons belonging to that assembly, producing one genome-level chromosome feature vector. The resulting genome-level chromosome vectors were subsequently mapped back to individual plasmids using Assembly_ID. When a genome contained multiple plasmids, each plasmid retained its own tRNA-presence or tRNA-copy-number response but received the same host-chromosome predictor vector. Thus, the prediction unit remained the individual plasmid rather than the host genome. LightGBM models with the same hyperparameters used in plasmid-based analyses were trained to predict plasmid tRNA presence and total tRNA abundance under identical genome-blocked partitions.

### Temporal holdout validation on an independent RefSeq cohort

An independent temporal validation cohort was constructed from RefSeq assemblies released between June 2025 and August 2026, comprising 10,772 GBFF assemblies that yielded 15,220 plasmids, including 1,111 tRNA-positive plasmids. Non-tRNA product counts were projected onto the fixed 29,267-feature vocabulary established from the original cohort. Features absent from this vocabulary were treated as out-of-vocabulary and excluded, with 98.11% of predictor instances successfully mapped to the predefined feature space. All tRNA-derived features were removed to prevent label leakage. The independent temporal cohort was used solely for final evaluation and was not involved in model training, feature selection, hyperparameter tuning, or threshold optimization.

### Feature importance and functional enrichment analyses

Feature importance was quantified from the optimized LightGBM models using gain-based importance scores^29^. To assess the extent to which predictive performance was concentrated within a limited subset of genes, models were retrained using the top-ranked features and compared with the full-feature model. Functional composition of the top-ranked features was summarized using curated KEGG/product-level categories^54,55^. KEGG categories were derived from a manually downloaded KEGG BRITE keyword workbook. Product names and category keywords were normalized by lowercasing and removing non-alphanumeric characters, and each product was assigned to all categories whose keyword patterns matched the normalized product name by case-insensitive regular-expression matching. This resulted in a multilabel mapping, with products lacking a matching keyword retained as unclassified.

Protein-coding sequences were also functionally annotated using eggNOG-mapper^32,33^, and corresponding COG/NOG assignments were linked to gene-product features used in the machine-learning framework. For genes associated with multiple COG categories, all non-ambiguous assignments were retained. COG categories were further grouped into higher-level functional classes for visualization and interpretation. To identify functional modules most strongly associated with plasmid tRNA repertoires, separate LightGBM models were trained using feature genes from individual COG categories. Products were assigned to COG categories using the multilabel mapping described above; products assigned to multiple categories were retained in each corresponding category. For each COG category, a separate LightGBM model was trained using only features assigned to that category, with predictive performance evaluated for both classification (mean test AUPRC across three independent 80:20 train–test splits) and regression (mean test R^2^ across the corresponding splits), allowing comparison of the relative contributions of different biological functions to plasmid tRNA organization.

### Calculation of codon compensation index (CCI)

To quantify the extent to which coding sequences preferentially use codons supported by plasmid-encoded tRNAs, we developed a codon compensation index (CCI). Codon usage can reflect translational selection and adaptation to available tRNA pools, but conventional codon-usage metrics quantify related yet distinct properties, such as overall synonymous codon bias or adaptation to a reference set of highly expressed genes^16–18^. CCI was therefore designed specifically to test whether codons decoded by plasmid-derived tRNAs are enriched relative to the expectation under equal synonymous codon usage. For each coding sequence, codon counts were calculated after excluding the terminal stop codon. Methionine and tryptophan were excluded because they lack synonymous codons.

Plasmid-derived translational supply was calculated from annotated plasmid tRNA anticodons, each mapped to its cognate codon by reverse complement. Wobble pairing was not considered^56^. For each amino acid, we compared the observed fraction of codons supported by plasmid-encoded tRNAs with the expected fraction under equal synonymous codon usage. CDS-level CCI was calculated as the mean enrichment across all informative amino-acid families:

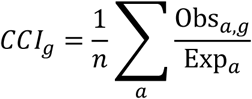

where Obs*_a_*_,*g*_ denotes the observed fraction of supported synonymous codons for amino acid *a* in gene *g*, and Exp*_a_* denotes the corresponding expectation under unbiased synonymous codon usage, and *n* is the number of informative synonymous amino-acid families in the CDS. A CCI greater than 1 indicates preferential use of codons decoded by plasmid-encoded tRNAs.

Unless otherwise specified, analyses used a pooled plasmid-tRNA definition, in which all plasmid-encoded tRNAs within a genome were combined into a single translational resource pool. This enabled direct comparison of chromosomal genes, genes on tRNA-positive plasmids, and genes on tRNA- negative plasmids within the same genomic background. In a complementary plasmid-specific analysis, CCI was recalculated for the CDSs of each tRNA-positive plasmid using only the tRNAs encoded by that same plasmid, thereby quantifying the extent to which its CDS codon usage relied on its own tRNA repertoire. Genes lacking informative synonymous codons were excluded.

For the primary comparison in Fig. 3B, CDS-level CCI values were used directly as independent observations. Analyses were restricted to genomes containing a chromosome, at least one tRNA- positive plasmid, and at least one tRNA-negative plasmid. Overall differences among the three categories (chromosomal genes, genes on tRNA-positive plasmids, and genes on tRNA-negative plasmids) were assessed using the Kruskal–Wallis test, followed by pairwise two-sided Mann–Whitney U tests with Benjamini–Hochberg correction for multiple comparisons.

### CCI aggregation and functional analyses

For phylogenetic comparisons, plasmid-level CCI was calculated by treating each plasmid as a single coding sequence, in which all coding regions on the plasmid were combined to quantify its overall codon compensation relative to the plasmid-borne tRNA repertoire. Plasmid-level CCI values were then summarized across phyla.

For product-level analyses, CCI values were aggregated across genes sharing the same annotated product descriptors and summarized using median CCI. Functional analyses were performed using the multilabel noS COG mapping described above, excluding function-unknown assignments. Products assigned to multiple COG categories were retained in each corresponding category, and CDS-level CCI summarized by COG category CCI values were summarized by their median.

### Central-dogma module annotation and prediction

To evaluate whether plasmid informational architecture extends beyond tRNAs, we constructed a project-specific central-dogma descriptor dictionary from the product qualifiers of PGAP-annotated features in the complete bacterial and archaeal RefSeq assemblies analyzed in this study. PGAP provided the standardized product vocabulary but did not assign central-dogma module labels. Candidate descriptors were retrieved using a high-sensitivity vocabulary covering process names, component names and recognized protein-family synonyms, including Rep-, Par-, Rec-, DNA polymerase, primase and helicase terms for replication; RNA polymerase, sigma-factor, Rho, Nus and transcriptional-regulator terms for transcription; and ribosomal protein, ribosome-biogenesis, aminoacyl-tRNA synthetase, rRNA, tRNA and translation-factor terms for translation. Every candidate descriptor was then manually reviewed for a direct mechanistic role in replication, transcription or translation and assigned to one of 14 mutually exclusive subcategories. Descriptors denoting hypothetical or uncharacterized proteins, or products without a sufficiently specific central-dogma function, were excluded.

The functional boundaries of the subcategories were defined from established reviews and reference resources. Replication comprised replication initiation^57^, replication machinery and elongation^58^, maintenance and partitioning^59^, and DNA repair and recombination^60^. Transcription comprised core RNA-polymerase machinery^61^, initiation and promoter recognition^62^, termination and antitermination^63^, and transcriptional regulation^64^. Translation comprised ribosomal proteins^65^, ribosome assembly and maturation^66^, aminoacyl-tRNA synthesis^67^, rRNAs, tRNAs, and translation factors and associated control processes^68^. Candidate keywords were used only to assemble the manually reviewed dictionary; plasmid genes were not classified using substring, regular-expression or fuzzy matching. Before exact matching, both curated and observed product strings were stripped of leading and trailing whitespace, consecutive internal whitespace was collapsed to a single space, and strings were compared case-insensitively. A curated entry was retained only when its normalized complete string exactly matched the Product field or corresponding internal feature code in the plasmid feature dictionary. Duplicate mappings were collapsed at the feature-code level. This procedure identified 1,386 unique target features: 402 replication, 611 transcription and 373 translation features.

Module predictability was evaluated using LightGBM classifiers and regressors. For each prediction task, features belonging to the target module were removed from the input matrix to prevent data leakage. Additional robustness analyses were performed by excluding all central-dogma-associated features or by restricting predictors to other central-dogma modules. Classification performance was assessed using AUPRC, whereas regression performance was evaluated using R². COG-defined J, K, and L categories were analyzed independently as annotation-based sensitivity controls.

## Data Availability

All the data associated with this work are available at the GitHub repository (https://github.com/zhongyuxuan11/Informational-architecture-organizes-plasmid-genomes).

## Code Availability

All codes are available at the GitHub repository (https://github.com/zhongyuxuan11/Informational-architecture-organizes-plasmid-genomes).

## Acknowledgement

This study was supported by the National Key R&D Program of China (2024YFA0920200 to TW), the National Natural Science Foundation of China (12401660 and 32470701 to TW), and the Shenzhen Institute of Synthetic Biology Scientific Research Program (HSE499011086 to TW). We are grateful to the Shenzhen Infrastructure for Synthetic Biology for providing instrument support and technical assistance.

