## Supplementary Figures for "Informational architecture organizes plasmid genomes"

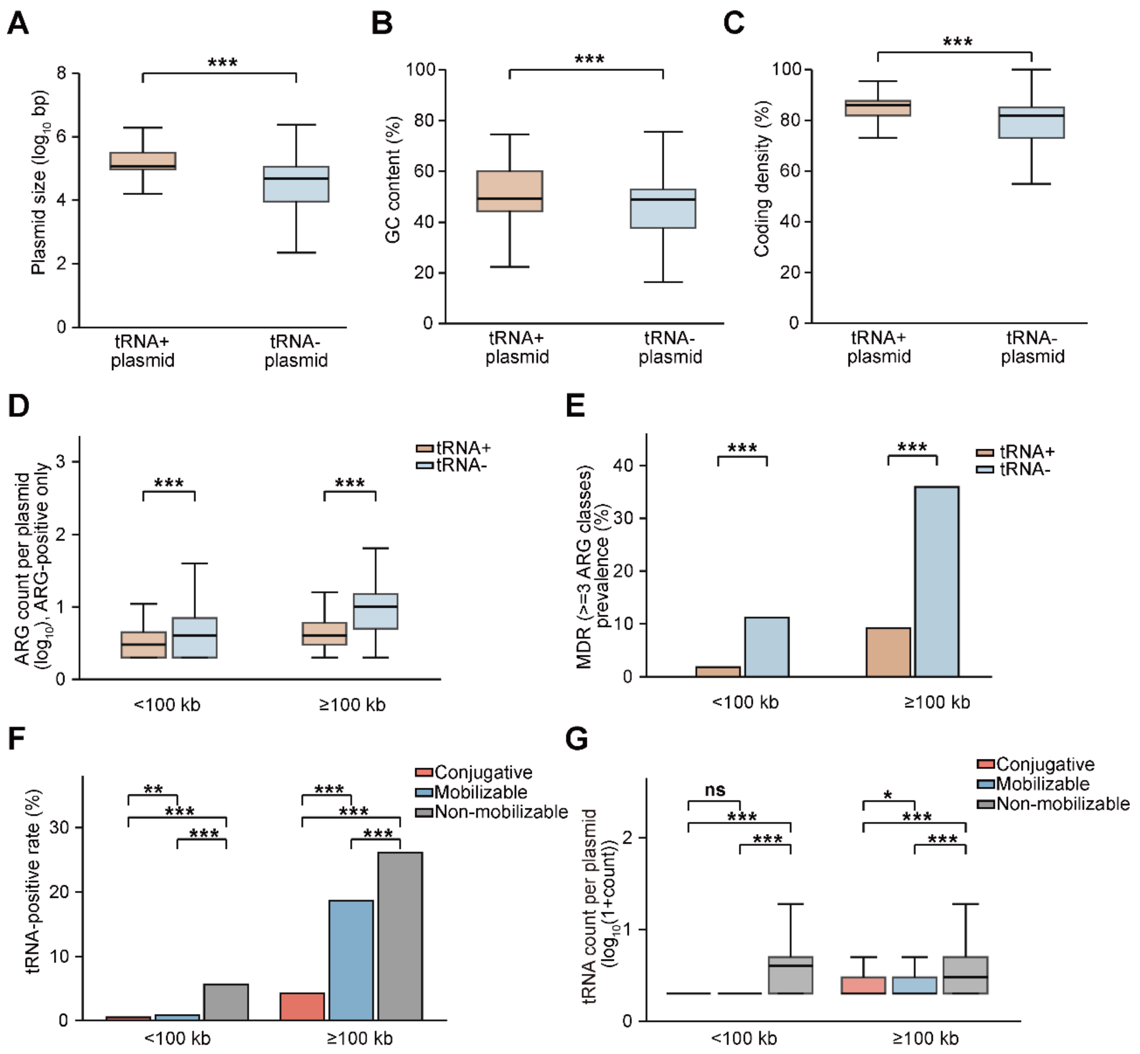


**Supplementary Figure S1| Basic genomic features of tRNA-encoding plasmids and size-stratified analyses of antibiotic resistance and mobility associations.**

(A–C) Comparison of plasmid size, GC content, and coding density between tRNA-positive and tRNA-negative plasmids. Groups were compared using two-sided Mann-Whitney U tests.

(D) Abundance of antibiotic resistance genes (ARGs) among ARG-positive plasmids, stratified by plasmid size (<100 kb vs. ≥100 kb). Within each size stratum, tRNA-positive and tRNA-negative plasmids were compared using a two-sided Mann-Whitney U test on raw counts; the two P values were adjusted using the Benjamini-Hochberg procedure.

(E) Prevalence of multidrug resistance (MDR) within the same size strata. Within each stratum, tRNA-positive and tRNA-negative plasmids were compared using a two-sided Fisher's exact test, followed by Benjamini-Hochberg adjustment across the two tests.

(F) Prevalence of tRNA-encoding plasmids across predicted mobility classes (Conjugative, Mobilizable, Non-mobilizable), shown separately for each size stratum. All three pairwise comparisons within each stratum were performed using two-sided Fisher's exact tests with Benjamini-Hochberg adjustment.

(G) tRNA gene counts per plasmid among tRNA-carrying plasmids, stratified by mobility class and size group. Within each stratum, the omnibus comparison used a Kruskal-Wallis test, followed by all three pairwise two-sided Mann-Whitney U tests with Benjamini-Hochberg adjustment. Boxes show the interquartile range (IQR) and median; whiskers extend to the most extreme values within 1.5 × IQR. In A and B, asterisks denote raw P values from single tests; in D-G, asterisks denote Benjamini-Hochberg-adjusted q values. ns, P or q >= 0.05; *P or q < 0.05; **P or q < 0.01; ***P or q < 0.001.

**A**

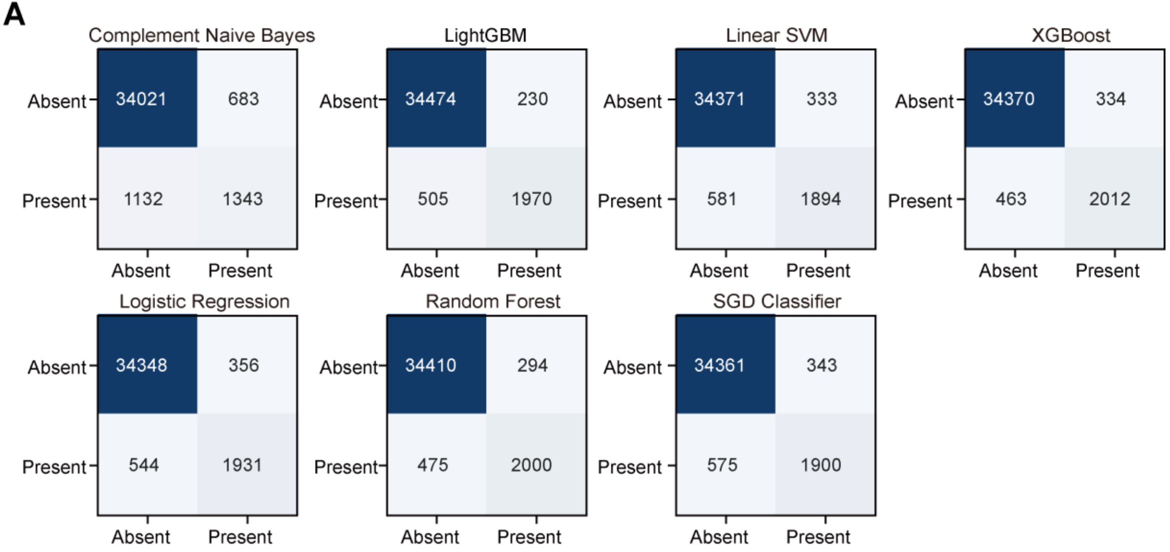


Mean ± SD, n = 3 independent 80:20 splits

| Model | AUPRC | AUROC | Precision | Recall | F1 |
|---|---|---|---|---|---|
| Random Forest | 0.906 ± 0.021 | 0.985 ± 0.004 | 0.872 ± 0.024 | 0.808 ± 0.030 | 0.839 ± 0.020 |
| LightGBM | 0.900 ± 0.015 | 0.979 ± 0.002 | 0.896 ± 0.030 | 0.796 ± 0.007 | 0.843 ± 0.017 |
| XGBoost | 0.897 ± 0.012 | 0.979 ± 0.002 | 0.859 ± 0.037 | 0.813 ± 0.003 | 0.835 ± 0.016 |
| Linear SVM | 0.865 ± 0.014 | 0.966 ± 0.005 | 0.851 ± 0.019 | 0.765 ± 0.020 | 0.806 ± 0.003 |
| Logistic Regression | 0.865 ± 0.014 | 0.969 ± 0.002 | 0.845 ± 0.022 | 0.780 ± 0.009 | 0.811 ± 0.009 |
| SGD Classifier | 0.864 ± 0.008 | 0.971 ± 0.003 | 0.847 ± 0.009 | 0.768 ± 0.018 | 0.805 ± 0.006 |
| Complement Naive Bayes | 0.506 ± 0.022 | 0.777 ± 0.008 | 0.665 ± 0.048 | 0.543 ± 0.001 | 0.597 ± 0.019 |

**B**

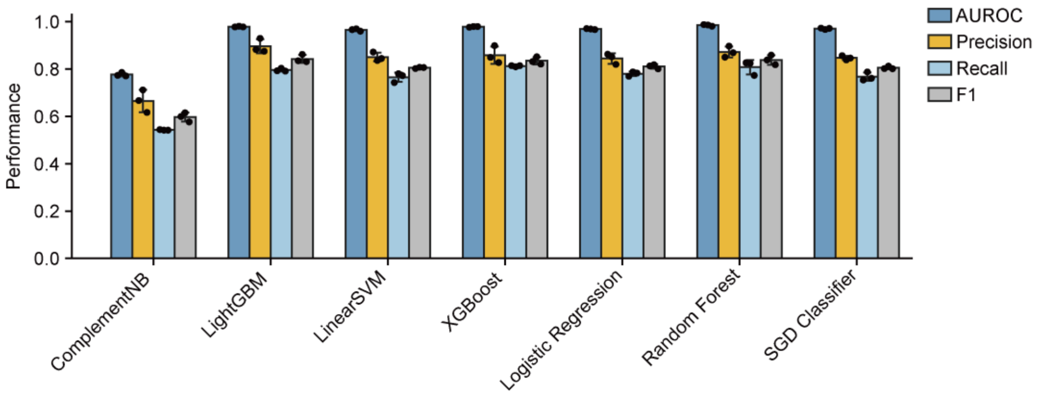


**Supplementary Figure S2| Complementary performance metrics for the seven-model tRNA-presence classifier benchmark.**

(A) Confusion matrices for the held-out test partitions across three independent random 80:20 train-test replicates, with the accompanying table summarizing held-out test-set AUPRC, AUROC, precision, recall and F1 as mean ± SD across the three replicates. All seven classifiers underwent hyperparameter optimization using 50 candidate configurations per classifier, evaluated exclusively on an internal validation subset of each training partition using AUPRC as the selection criterion.

(B) Held-out test-set AUROC, precision, recall and F1 for each model. Bars show mean test performance, error bars show SD, and black points show raw replicate values for seeds 42, 43 and 44.

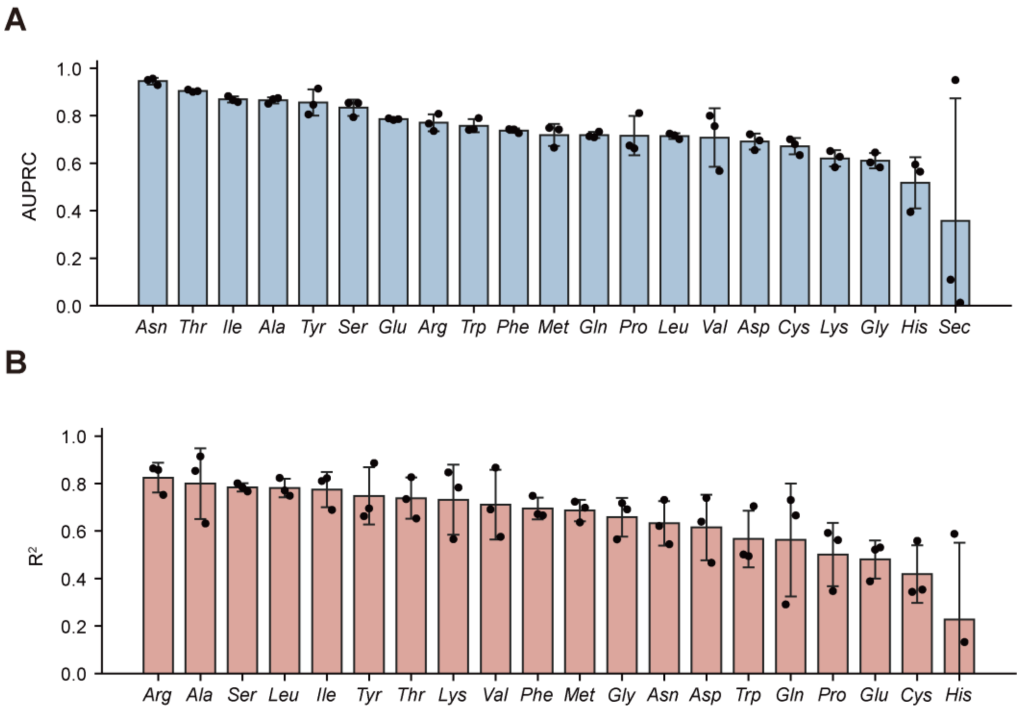


**Supplementary Figure S3 | Predictability of individual plasmid-encoded tRNA types.**

(A) LightGBM classification performance for predicting the presence or absence of each of the 21 plasmid-encoded tRNA types.

(B) LightGBM regression performance for predicting raw copy number among plasmids positive for the corresponding tRNA type. tRNA-Sec regression was omitted because the positive-test target was constant and $R^2$ was undefined. Bars show the mean across three random 80:20 train-test replicates, error bars show SD, and black points show the raw replicate values.

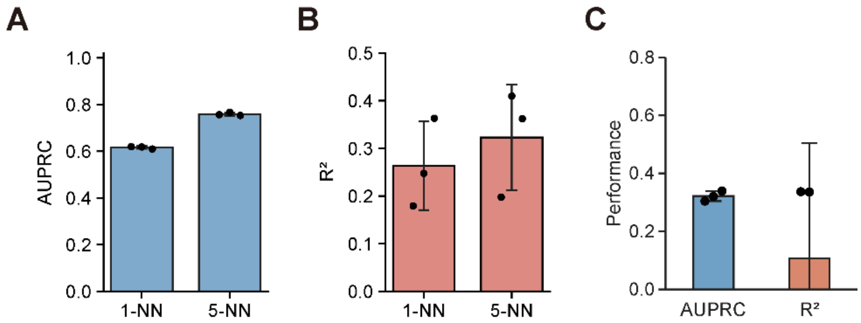


**Supplementary Figure S4 | Alternative baselines and external validation**.

(A) Classification AUPRC of 1-nearest-neighbor and 5-nearest-neighbor models based on plasmid gene-content similarity.

(B) Regression $R^2$ of the corresponding nearest-neighbor baselines for raw plasmid tRNA abundance.

(C) Host-chromosome baseline. LightGBM models were trained using chromosome-derived non-tRNA product-count vectors mapped to the same target plasmids, labels, splits and evaluation protocol as the plasmid-product models; plasmid products were not used as predictors.

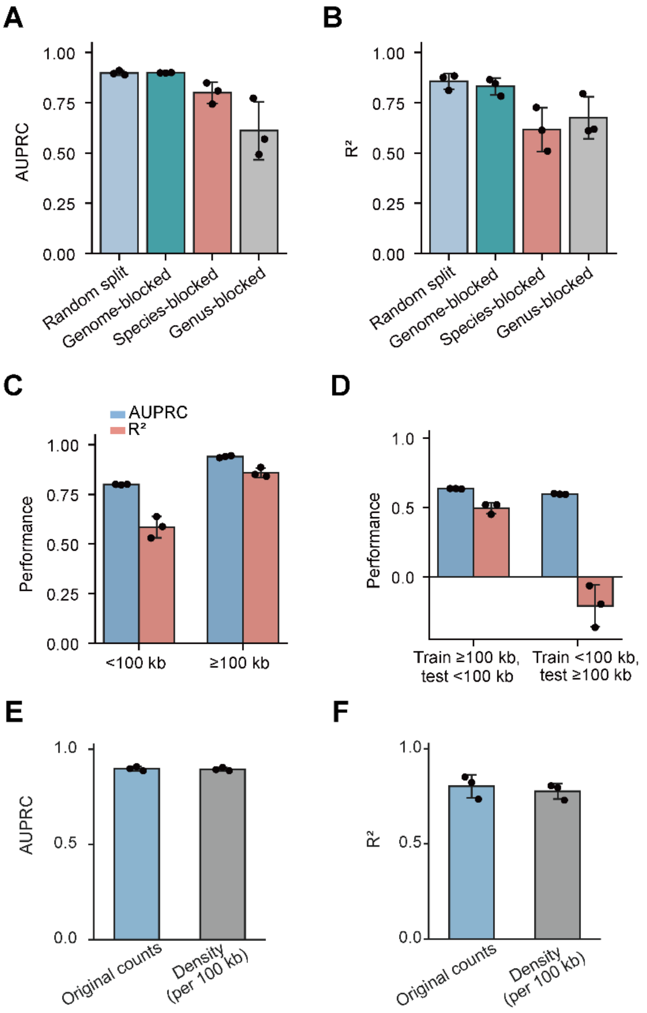


**Supplementary Figure S5 | Robustness of LightGBM prediction across split strategies and plasmid-size strata.**

(A) Classification AUPRC under random, genome-blocked, species-blocked and genus-blocked split strategies.

(B) Regression $R^2$ under the same split strategies.

(C) Within-size-bin prediction performance for <100 kb and ≥100 kb plasmids.

(D) Reciprocal size-transfer performance when models trained in one plasmid-size class were tested on the other class; both classification AUPRC and regression $R^2$ are shown. For random and within-bin analyses, bars show means from three 80:20 replicates, error bars show SD, and black points show raw replicate values. In size-transfer analyses, source and target size classes were fixed and seeds varied model stochasticity.

(E, F) Size-normalization sensitivity analysis. Model performance was compared between two feature sets: original model (absolute product counts) and density model (product abundance per 100 kb, normalized by plasmid length). Classification and regression used identical train–test splits (seeds 42–44) and locked LightGBM parameters. Bars/points indicate mean ± SD across three splits.

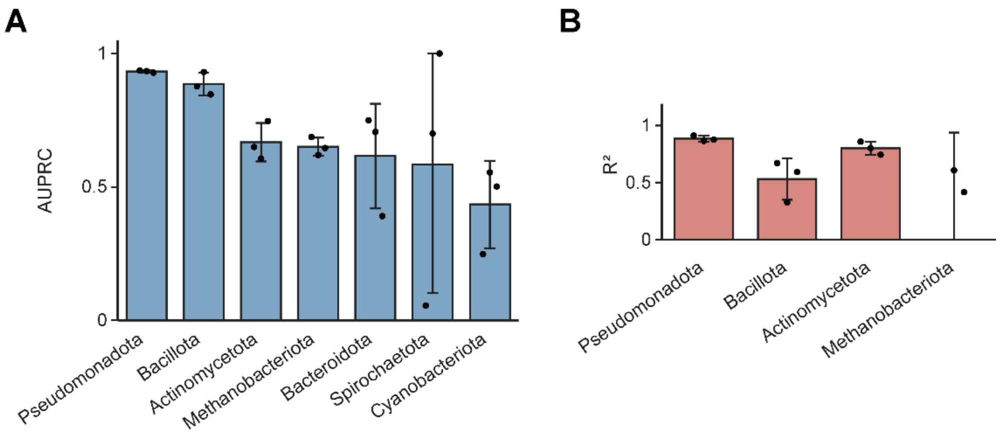


**Supplementary Figure S6 | Phylum-stratified prediction performance.**

(A) Classification AUPRC and (B) regression $R^2$ were calculated within phyla after fitting the single locked model for each replicate; models were not retrained within phyla. Phyla were plotted only when the corresponding test subset had more than 10 samples and valid targets. Classification panels required both tRNA-positive and tRNA-negative plasmids within the phylum-specific test subset; regression panels used tRNA-positive plasmids and required nonconstant tRNA counts. Bars show mean performance across the three held-out test replicates, error bars show SD, and black points show raw replicate values.

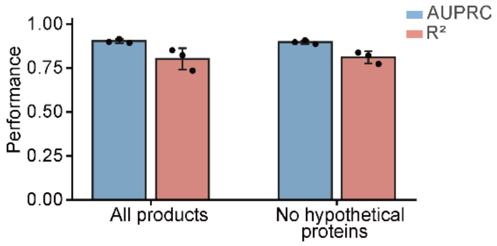


**Supplementary Figure S7 | Effect of removing hypothetical protein features.** Classification AUPRC and regression $R^2$ were compared between the full non-tRNA product matrix and the no-hypothetical feature matrix after removing the standardized hypothetical-protein feature. Models used the same locked LightGBM hyperparameters and matched split manifests. Bars show the mean across three random 80:20 train-test replicates, error bars show SD, and black points show raw replicate values.

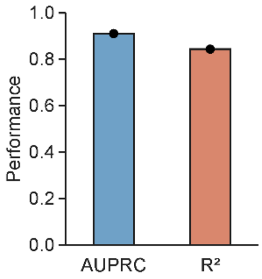


**Supplementary Figure S8 | Temporal validation of plasmid tRNA prediction.** A LightGBM classification model and a LightGBM regression model were trained once on the complete original RefSeq plasmid cohort using the fixed historical feature vocabulary, then evaluated without retraining on an independent non-overlapping later RefSeq plasmid cohort. Black points indicate the single external-validation estimates.

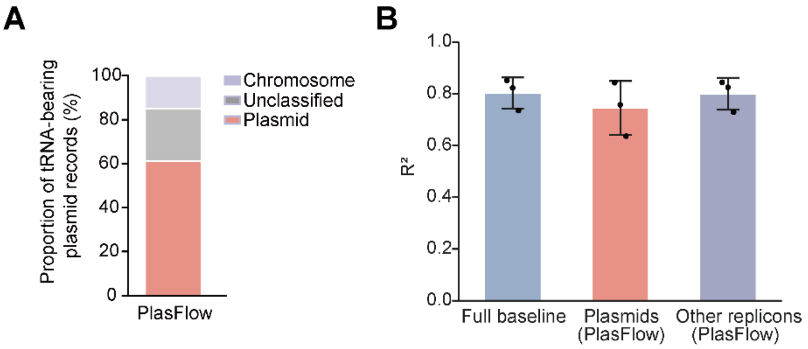


**Supplementary Figure S9 | PlasFlow-based replicon annotation sensitivity analysis.**

(A) PlasFlow assignments for tRNA-bearing plasmid records, classified as plasmid-like, chromosome-like or unclassified.

(B) Regression $R^2$ for the full held-out baseline set, the subset classified as plasmid-like by PlasFlow, and other replicons (chromosome-like plus unclassified/non-plasmid PlasFlow assignments). Locked LightGBM regression model was refitted on each replicate's development partition before held-out prediction; the PlasFlow sensitivity step then recalculated metrics on test-set subsets without additional retraining or feature modification. Bars show mean $R^2$ across three random 80:20 replicates, error bars show standard deviation, and black points show raw replicate values.

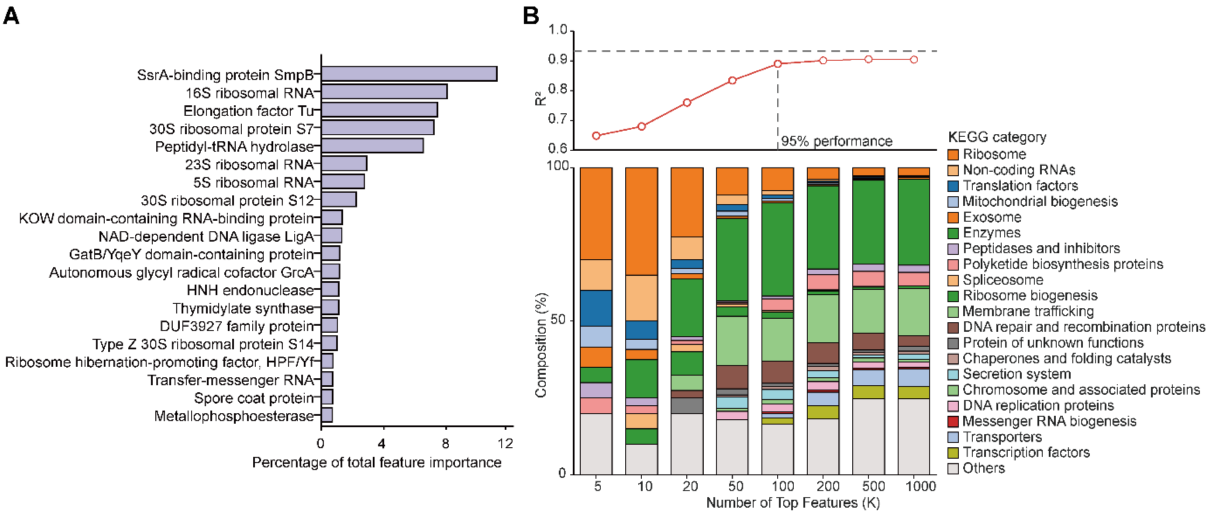


**Supplementary Figure S10 | Regression-model feature importance and Top-K feature sufficiency.**

(A) Top-ranked predictors from the LightGBM regression model for plasmid tRNA abundance, shown as percentage of total normalized gain.

(B) Regression $R^2$ after retraining models using increasing numbers of top-ranked features, together with the functional-category composition of the corresponding top-ranked features. Feature-importance and Top-K analyses used the no-hypothetical feature matrix. Performance curves show means across three random 80:20 train-test replicates, error bars show SD, and black points show raw replicate values where plotted. The dashed line indicates 95% of full-model regression performance.

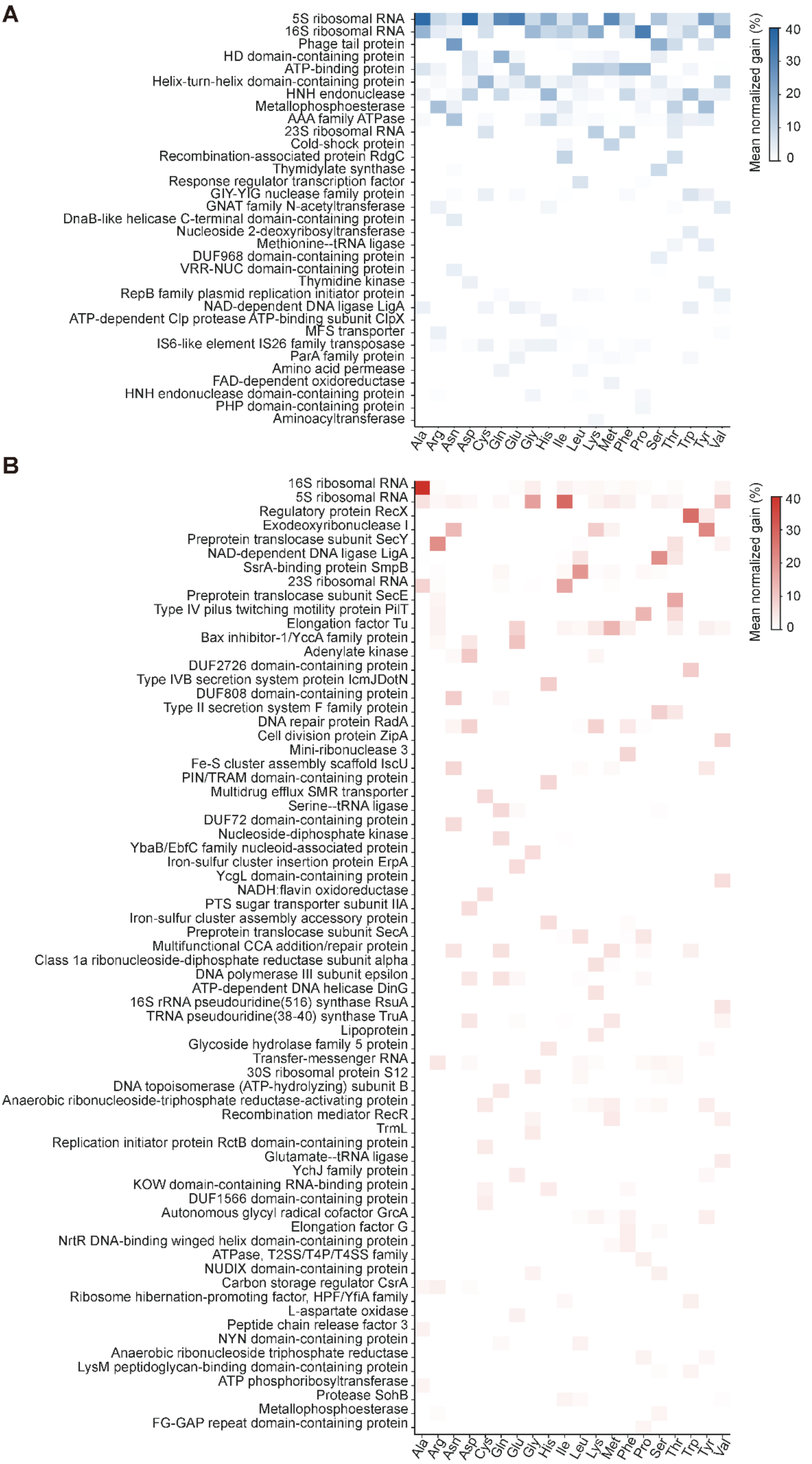
A
5S ribosomal RNA
16S ribosomal RNA
Phage tail protein
HD domain-containing protein
ATP-binding protein
Helix-turn-helix domain-containing protein
HNH endonuclease
Metallophosphoesterase
AAA family ATPase
23S ribosomal RNA
Cold-shock protein
Recombination-associated protein RdgC
Thymidylate synthase
Response regulator transcription factor
GIY-YIG nuclease family protein
GNAT family N-acetyltransferase
DnaB-like helicase C-terminal domain-containing protein
Nucleoside 2-deoxyribosyltransferase
Methionine--tRNA ligase
DUF968 domain-containing protein
VRR-NUC domain-containing protein
Thymidine kinase
RepB family plasmid replication initiator protein
NAD-dependent DNA ligase LigA
ATP-dependent Clp protease ATP-binding subunit ClpX
MFS transporter
IS6-like element IS26 family transposase
ParA family protein
Amino acid permease
FAD-dependent oxidoreductase
HNH endonuclease domain-containing protein
PHP domain-containing protein
Aminoacyltransferase
Ala Arg Asn Asp Cys Gln Glu Gly His Ile Leu Lys Met Phe Pro Ser Thr Trp Tyr Val
Mean normalized gain (%)
40
30
20
10
0
B
16S ribosomal RNA
5S ribosomal RNA
Regulatory protein RecX
Exodeoxyribonuclease I
Preprotein translocase subunit SecY
NAD-dependent DNA ligase LigA
SsrA-binding protein SmpB
23S ribosomal RNA
Preprotein translocase subunit SecE
Type IV pilus twitching motility protein PilT
Elongation factor Tu
Bax inhibitor-1/YccA family protein
Adenylate kinase
DUF2726 domain-containing protein
Type IVB secretion system protein IcmJDotN
DUF808 domain-containing protein
Type II secretion system F family protein
DNA repair protein RadA
Cell division protein ZipA
Mini-ribonuclease 3
Fe-S cluster assembly scaffold IscU
PIN/TRAM domain-containing protein
Multidrug efflux SMR transporter
Serine--tRNA ligase
DUF72 domain-containing protein
Nucleoside-diphosphate kinase
YbaB/EbfC family nucleoid-associated protein
Iron-sulfur cluster insertion protein ErpA
YcgL domain-containing protein
NADH:flavin oxidoreductase
PTS sugar transporter subunit IIA
Iron-sulfur cluster assembly accessory protein
Preprotein translocase subunit SecA
Multifunctional CCA addition/repair protein
Class 1a ribonucleoside-diphosphate reductase subunit alpha
DNA polymerase III subunit epsilon
ATP-dependent DNA helicase DinG
16S rRNA pseudouridine(516) synthase RsuA
TRNA pseudouridine(38-40) synthase TruA
Lipoprotein
Glycoside hydrolase family 5 protein
Transfer-messenger RNA
30S ribosomal protein S12
DNA topoisomerase (ATP-hydrolyzing) subunit B
Anaerobic ribonucleoside-triphosphate reductase-activating protein
Recombination mediator RecR
TrmL
Replication initiator protein RctB domain-containing protein
Glutamate--tRNA ligase
YchJ family protein
KOW domain-containing RNA-binding protein
DUF1566 domain-containing protein
Autonomous glycyl radical cofactor GrcA
Elongation factor G
NrtR DNA-binding winged helix domain-containing protein
ATPase, T2SS/T4P/T4SS family
NUDIX domain-containing protein
Carbon storage regulator CsrA
Ribosome hibernation-promoting factor, HPF/YfiA family
L-aspartate oxidase
Peptide chain release factor 3
NYN domain-containing protein
Anaerobic ribonucleoside triphosphate reductase
LysM peptidoglycan-binding domain-containing protein
ATP phosphoribosyltransferase
Protease SohB
Metallophosphoesterase
FG-GAP repeat domain-containing protein
Ala Arg Asn Asp Cys Gln Glu Gly His Ile Leu Lys Met Phe Pro Ser Thr Trp Tyr Val
Mean normalized gain (%)
40
30
20
10
0

**Supplementary Figure S11 | Type-specific feature importance for plasmid tRNA prediction.** (A) Gain-based feature-importance profiles for type-specific classification models predicting the presence or absence of each of the 20 plasmid-encoded tRNA types. (B) Gain-based feature-importance profiles for type-specific regression models predicting tRNA copy number for each tRNA type. Models used the no-hypothetical feature matrix and locked LightGBM hyperparameters. Feature-importance values were normalized as percentages of total gain within each fitted model. Classification and regression plots for the same tRNA type use the union of their highest-ranking features to facilitate direct comparison; tRNA-Sec regression was excluded because the regression target was constant and $R^2$ was undefined.

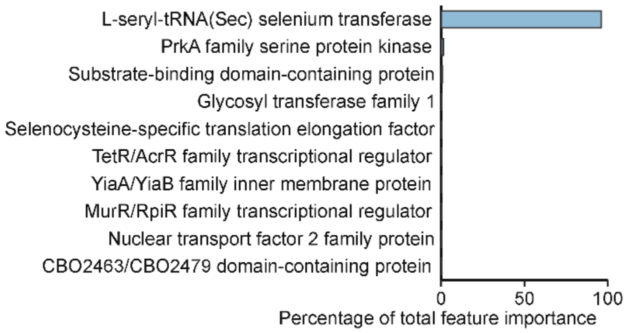


**Supplementary Figure S12 | Feature importance for plasmid tRNA-Sec prediction.** Gain-based feature importance from the LightGBM classification model for predicting the presence or absence of plasmid-encoded tRNA-Sec. Features are shown as percentages of total normalized gain for the classification model evaluated by AUPRC.

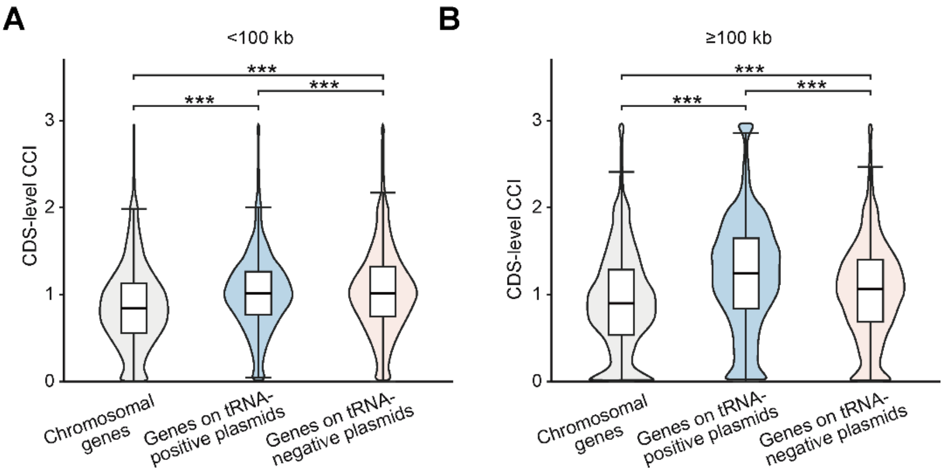


**Supplementary Figure S13 | Size-stratified analyses of codon compensation index (CCI) in relation to plasmid-encoded tRNAs and CDS-level CCI.**

(A, B) CDS-level CCI distributions for chromosomal, tRNA+ plasmid, and tRNA− plasmid genes, separately for plasmids <100 kb and ≥100 kb. CDS-level CCI is the CCI of each individual valid CDS, calculated using the genome-pooled plasmid-tRNA repertoire. Global differences were assessed by Kruskal–Wallis test followed by pairwise two-sided Mann–Whitney U tests with Benjamini–Hochberg correction. *$q < 0.05$; **$q < 0.01$; ***$q < 0.001$.

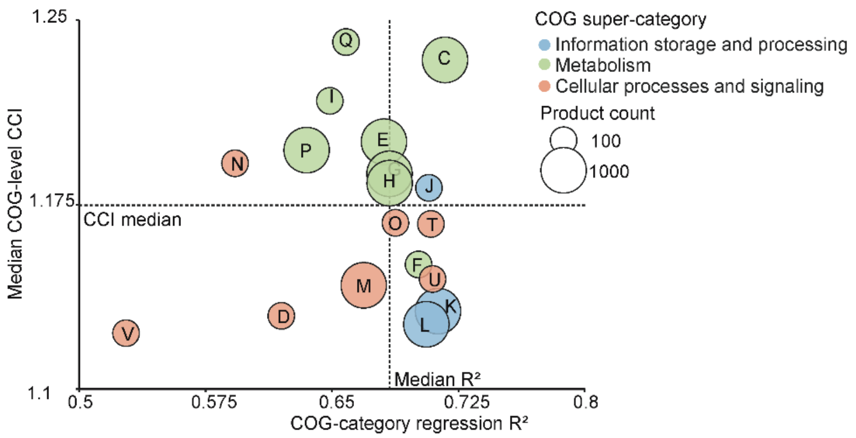


**Supplementary Figure S14 | Relationship between COG-category median product-level CCI and regression performance of COG-restricted models.** The x-axis shows the mean regression $R^2$ of LightGBM models trained using only features assigned to that COG category, and the y-axis shows the median product-level CCI of products assigned to the same category. Each point represents a COG category; colors indicate functional superclass. Dashed lines mark median values.

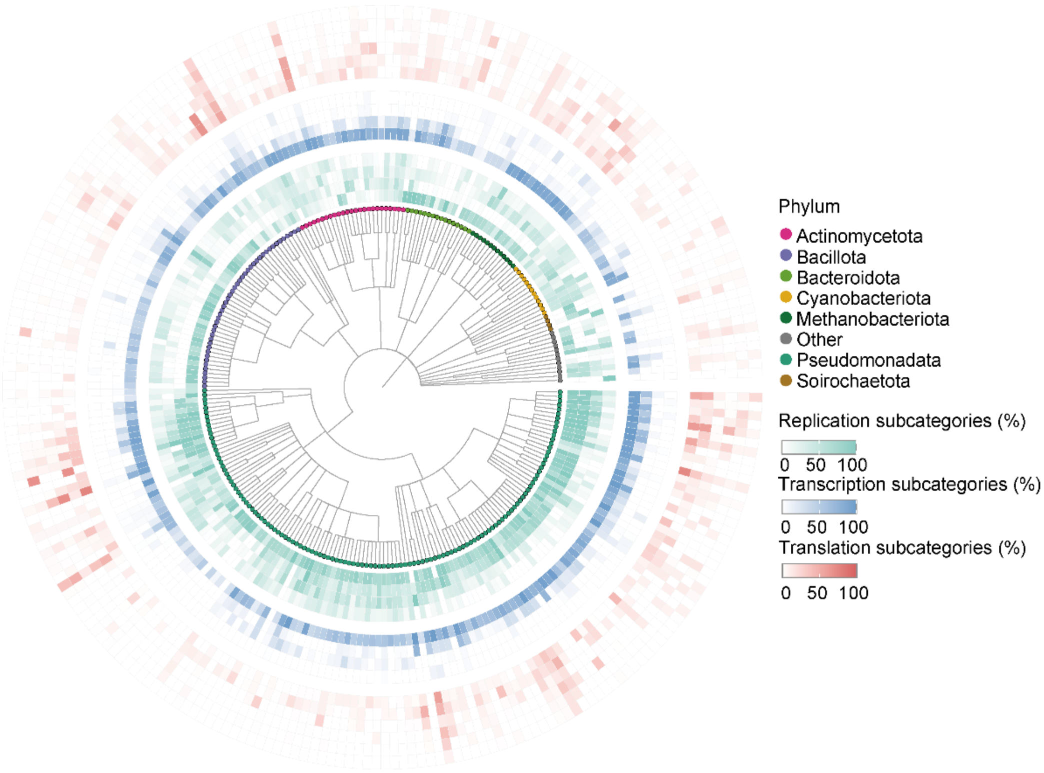


**Supplementary Figure S15 | Landscape of central dogma–associated gene categories across prokaryotic genera.** Radial phylogenetic heatmap showing the prevalence of replication-, transcription-, and translation-associated gene subcategories across plasmid-carrying genera. Tree-tip colors indicate different phyla. The 14 concentric rings correspond sequentially to four replication subcategories (replication initiator, maintenance/partition, DNA repair and recombination, and machinery/elongation), four transcription subcategories (transcription regulation, initiation and promoter recognition, termination and anti-termination, and core transcription machinery) and six translation subcategories (tRNA, regulation and control, aminoacyl-tRNA synthetases, ribosomal RNA, ribosomal proteins, and ribosome assembly and maturation). Color intensity indicates the prevalence of each subcategory within each genus, using separate color scales for replication, transcription, and translation.

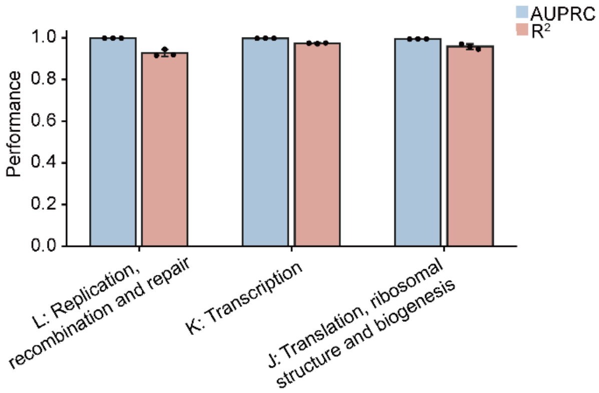


**Supplementary Figure S16 | COG functional categories L, K, and J**. For each task, all features assigned to the corresponding target COG category were removed from the predictor matrix before model training, and the remaining non-target plasmid features were used as predictors. Classification was evaluated by AUPRC and regression by $R^2$ across three target-stratified random 80:20 train-test replicates. Bars show replicate means, error bars show SD, and black points show raw replicate values. Positive-class prevalence should be considered when interpreting AUPRC because L, K and J are common categories in the test sets.
